# Neural responses to instructed positive couple interaction: An fMRI study on compliment sharing

**DOI:** 10.1101/2022.06.15.496238

**Authors:** Monika Eckstein, Gabriela Stößel, Martin Fungisai Gerchen, Edda Bilek, Peter Kirsch, Beate Ditzen

## Abstract

Love is probably the most fascinating feeling that a person ever experiences. However, little is known about what is happening in the brains of a romantic couple –the central and most salient relationship during adult age– while they are particularly tender and exchanging loving words with one another.

To gain insight into nearly natural couple interaction, we collected data from N=84 individuals (including N=43 heterosexual couples) simultaneously in two functional magnetic resonance imaging scanners, while they sent and received compliments, i.e. short messages about what they liked about each other and their relationship. Activation patterns during compliment sharing in the individuals revealed a broad pattern of activated brain areas known to be involved in empathy and reward processing. Notably, the ventral striatum, including parts of the putamen, was activated particularly when selecting messages for the partner. This provides initial evidence that giving a verbal treat to a romantic partner seems to involve neural reward circuitry in the basal ganglia.

These results can have important implications for the neurobiological mechanisms protecting and stabilizing romantic relationships, which build a highly relevant aspect of human life and health.

## Introduction

In almost all human cultures, romantic love is viewed as a central concept to giving meaning and joy to a person’s life. Social identity theory states that individuals derive parts of their identity from belonging to a group, a family, or a romantic relationship (Scheepers and Ellemers, 2019) and such social identification is related to less harmful stress as mediated by social support (Haslam *et al*., 2005). Specifically being in a functional couple relationship is even linked to better health and longer lives (Braithwaite and Holt-Lunstad, 2017).

The health-related impact of couple relationships is very likely centrally mediated with interacting neural networks and structures, such as the limbic reward system and its neurotransmitters. An interaction of the neuromodulator oxytocin and the neurotransmitter serotonin in the nucleus accumbens (Dölen *et al*., 2013) has been shown to mediate social reward. Oxytocin interacting with dopamine has been suggested to contribute to the formation and maintenance of social bonds in animals (Bosch and Young, 2018) and in humans for instance via a positive evaluation of the own relationship (Scheele *et al*., 2013; Aguilar-Raab *et al*., 2019).

As parts of the dopaminergic reward system, the nucleus accumbens, together with putamen and ventral tegmental area (VTA) among others, are involved in the initiation of joyful behaviors and feelings in general, but especially social reinforcements (Izuma *et al*., 2008; Dölen *et al*., 2013). Functional activation of the dopaminergic reward systems might thus be one (of several) underlying mechanism supporting initializing and maintaining human couple relationships (Bartels & Zeki, 2004) and is therefore in the focus of the present study. In previous research, interacting with the partner or observing a partner picture was associated with elevated activation in the VTA, hippocampus, insula (Bartels and Zeki, 2004), anterior cingulate cortex (ACC) (Aron *et al*., 2005) posterior superior temporal sulcus (pSTS), and anterior temporal lobe (ATP)(Van der Gaag *et al*., 2007).

One precondition for functional social interaction and for romantic couple relationships in particular is a theory of mind (ToM), the ability to infer the status of knowledge of another person. ToM is related to activation of the superior temporal brain, temporal, and frontal areas (Dodell-Feder *et al*., 2015), while actual empathy recruits the anterior insula (Kennedy and Adolphs, 2012; Thornton *et al*., 2019). During empathy-related processes, the accumbens is also interacting with the ACC (Smith *et al*., 2021). In addition, the mirror neuron system, which comprises parietal and frontolateral brain areas, is involved in social perception and action (Mier *et al*., 2010).

Furthermore, social integration and the perception of belonging increase positive affect and self-esteem (Ellemers *et al*., 1999). Presumably, positive feedback acts as an indicator of social integration. For romantic couples, a constructive way of communication has been shown to be related to better relationship satisfaction and even to buffer a lack of sexual satisfaction (Litzinger and Gordon, 2005) and communicating compliments in everyday life has been linked to better relationship satisfaction with a stronger sensitivity towards compliments specifically in women (Doohan and Manusov, 2004). Imaging studies have shown that receiving compliments from a stranger or from one’s own mother involved the dorsolateral prefrontal cortex (DLPFC) (Hooley *et al*., 2005), ACC, and temporal areas (Miedl *et al*., 2016). Based on this, receiving compliments from the partner can be considered highly relevant to evaluating the social self, the level of integration and affection and, thereby, act in a particularly rewarding and health-beneficial way. To investigate the neural responses to tender partner compliments, we have adapted the previously established standard instructed partnership appreciation task (Pfeifer *et al*., 2020; Warth *et al*., 2020) for a functional imaging (fMRI) paradigm. We compared compliments from the partner to “self compliments” (i.e., attributes that the participants defined about themselves), since the mental reflection of positive attributes per se could improve mood (Nicolson *et al*., 2020) by activating reward-related brain areas (Izuma *et al*., 2008; Frewen *et al*., 2020).

For general compliment processing we expected that receiving compliments from the partner would result in elevated activation in a broad network including VTA, hippocampus, insula, ACC, and pSTS, (Van der Gaag *et al*., 2007). In addition, reward-related task phases as well as phases of reward anticipation (Filimon *et al*., 2020) should be related to activation in the dopaminergic system: the ventral striatum including the nucleus accumbens. While a participant is actively choosing a compliment, we expected activity known for reading and decision making, and when a participant is sending the compliment, the areas relevant for ToM should be activated, along with mirror neuron areas when they are observing partners’ reactions. These activation patterns should become evident using whole brain approaches.

## Methods

### Participants

Eighty six heterosexual participants (from 43 romantic couples) who were in love and exclusively dating for at least six months were recruited in the Rhine-Neckar metropolitan area, Germany; see Table 1 for sample characteristics. In addition to sociodemographic data, participants provided information on their relationship quality (Partnership Questionnaire, PFB (Hahlweg, 1979), including the scales for tenderness, conflict, and joint interests.

**Table 1.**
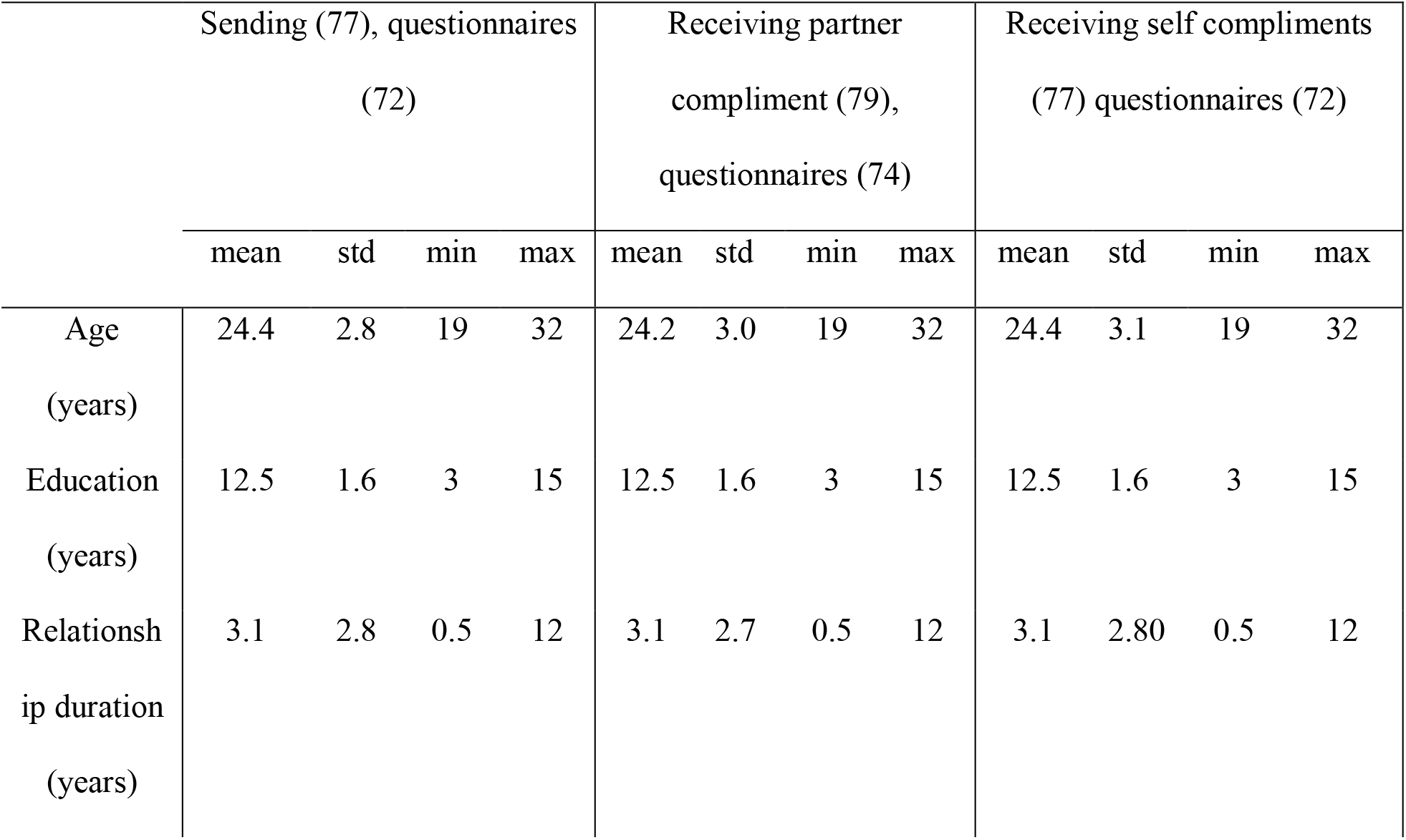
Sample characteristics. Available data presented for participants included for the corresponding fMRI analyses (number of participants in brackets):

Particularly happy couples (reporting at least 5 on a 6-point single-item rating scale on general relationship satisfaction) were included in the study. All participants were eligible for MRI scanning, right-handed, without history of mental disorders and knew sufficient German language to fully understand all instructions. Couples provided written informed consent and were reimbursed 80€ per couple for their participation. The study was conducted in accordance with the Declaration of Helsinki and approved by the Ethics Committee of Heidelberg University Medical Faculty (#2011-222N-MA).

### Paradigms

In an interview session with the individual participants prior to the MRI session, all participants were handed a list of 23 areas of individual traits and relationship aspects, based on factors of the PFB (e.g., trust, humor, intimacy). Based on these areas, participants were asked to generate up to 18 short positive messages (compliments) about their partner for use in the upcoming experiment. In addition, participants created up to 18 compliments about themselves to be viewed as control stimuli. The compliments were kept confidential until the MRI session. Non-German native speaking couples were allowed to provide compliments in their native language. The paradigm consisted of 15 trials per condition (receiving, sending, self compliment). In the send and receive compliment condition, each trial consisted of two phases, lasting 10s each. In the first phase, the sender chose one of four compliments shown on his/her screen, and the receiver waited for the partner to select the message. In the second phase of the trial, the compliment was revealed to both partners. In the self compliment paradigm, running on both scanners simultaneously, trials consisted of two phases as well: in the first phase, the text ‘Please wait, computer is choosing your compliment’ was displayed to both participants, in the second one, the text appeared: ‘Computer has chosen: compliment_text’. Both phases of each trial were jittered on average by 775ms, one whole trial lasted 32.5s. All texts were presented on the left-hand side of the screen. On the right-hand side, a live video of the partner taken with a wide-angle camera (MRC Systems GmbH, Heidelberg, Germany) in infrared light was shown continually during all paradigms in order to keep general effects of partner contact constant over conditions. The participants were randomly assigned to one of the two scanners. The order of which partner sent first, as well as the assignment of sexes to the scanner and the orders to the scanner were balanced. The temporal order of paradigms (partner vs. self) and the initial sender-sex-scanner matching was randomized and balanced across the sample. However, the first send/receive condition was always followed by the complimentary send/receive condition. The task followed an anatomical measurement and a joint attention paradigm (Bilek *et al*., 2015).

A follow-up questionnaire directly after the fMRI session assessed the participants’ overall evaluation of sending and receiving partner-compliments with: “How much did you enjoy sending and receiving the messages?” and “How much did you enjoy reading your own positive attributes?” on a 9-point scale from “not at all” to “very much”.

### Data acquisition

Data was acquired with two synchronized 3 Tesla Siemens Tim TRIO scanners, where one scanner was triggered by the other one. Twelve-channel head coils were used. A T2* gradient echo-planar imaging sequence was applied with the following parameters: 28 axial slices, with transversal orientation, oriented first to AC/PC line and then flipped by -25°, 4 mm slice thickness, 1 mm gap, field of view 192 mm, voxel size 3×3×4 mm3. Repetition time (TR) was 1.55s with sampling delay of 10ms, and 1.54s. Echo time was 30ms, flip angle 73°. Slices were acquired in descending order, with A/P phase encoding direction. The GRAPPA method with an acceleration factor of 2 was used. A total of 327 (triggering)/ 324 (triggered scanner) scans were collected per condition. The first 7 (triggering)/4 (triggered scanner) scans were discarded during conversion of DICOM-files into 4d niftis by MRIConvert (version. 2.0 rev. 216) to account for saturation effects, resulting in 320 scans available for analysis per condition.

A high-resolution (voxel size 1×1×1mm) T1 anatomical scan was acquired for individual anatomical registration purposes.

### Data analyses and preprocessing

fMRI data were analyzed using SPM12 (v771). The anatomical image was segmented and normalized to the SPM12 TPM MNI template. Preprocessing of the functional data involved slice-time correction, realignment to the mean image, and co-registration of the functional images (mean and others) to the anatomical image. The co-registered functional data was normalized to MNI space, resampled to 3mm^3^ voxels, and smoothed with a Gaussian kernel with full-width-at-half-maximum FWHM=8×8×8 mm. Volumes affected by small movement artefacts were identified with the ART toolbox (http://www.nitrc.org/projects/artifact_detect ; parameters: framewise displacement >0.5mm, image intensity change z>4, and exclusion criterion for a measurement: >25% affected volumes).

Of the original 86 fMRI measurements, we had to exclude nine from the activation analysis of the send-paradigm (resulting in N=77) and seven measurements from the activation analysis of the receive-paradigm (resulting in N=79) due to excessive head motion, technical problems, or aborted measurements due to time constraints. In total, this resulted in 14 participants having to be excluded from the comparison of the receive-paradigm with the self compliment-paradigm (N=72).

First, we analyzed the task-related activation in the individuals’ brains by means of general linear modeling. A first-level model with three sessions for the three separate conditions of the experiment was set up to allow for both within-session and across-session contrasts. With the conditions, the individual phases (waiting for and receiving a compliment, as well as selecting and observing shared compliments) were modeled as blocks. Signals from cerebrospinal fluid and white matter, 24 movement parameters (six standard parameters, their backward derivatives, and their squared versions), and ART dummy regressors were included as nuisance regressors. A high-pass filter with a frequency cutoff of 128s was applied, as well first-degree autoregression.

In the group analyses, age, sex and scanner were included as covariates. Analyses were conducted using one-sample t-tests over the respective contrasts. Contrasts of interests were [Receiving > Waiting] within blocks (partner compliment and self compliment) and [Receiving > Waiting] compared between blocks (partner compliment and self compliment) as well as a contrast between the active block [Choosing compliment > Observing sent compliment] and the passive block [Receiving > Waiting]. All activation results are reported with p<0.05 whole-brain FWE-corrected significance. Beta estimates were additionally extracted, only for visualization of the activity of the ventral striatum (anatomical region-of-interest from the AAL-90 atlas) during conditions (Fig. 4).

Questionnaire data was analyzed using SPSS 27 (IBM).

## Results

### Activation of individuals when receiving a compliment

When participants were passively receiving compliments (both from the partner and self compliments) as compared to the waiting phases (within blocks), increased activation in a broad network of IFG, DLPFC, VMPFC, midbrain-structures and temporal gyri was observed, see Tables 2 and 3 and Fig. 1a and b.

**Table 2.** Brain responses to (receiving partner compliments > waiting for partner compliments)

| peak | peak |  |  |  |  |
| --- | --- | --- | --- | --- | --- |
| p(FWE-corr) | T | x | y | z | Automated anatomical labeling atlas<br>(Tzourio-Mazoyer *et al.*, 2002) |
| < 0.001 | 16.79296303 | 3 | 14 | 65 | Superior frontal gyrus/Supplementary motor area |
| < 0.001 | 12.37275314 | -6 | 5 | 77 | Superior frontal gyrus |
| < 0.001 | 11.77529621 | 6 | 17 | 38 | Middle cingulate gyrus |
| < 0.001 | 15.70023251 | -39 | -70 | -22 | Cerebellum |
| < 0.001 | 15.21589851 | -42 | 17 | -28 | Superior temporal gyrus |
| < 0.001 | 15.20484543 | -45 | 17 | -7 | Inferior frontal gyrus |
| 0.0049 | 5.602838039 | 66 | -40 | 26 | Inferior parietal lobule/Supramarginal gyrus |
| 0.0065 | 5.522605896 | 66 | -58 | 11 | Superior temporal gyrus |
| 0.0447 | 4.955034733 | -15 | -25 | -31 | Cerebellum |

*Whole brain analyses; FWE corr, familywise error correction (applies to all tables)*

**Table 3.**
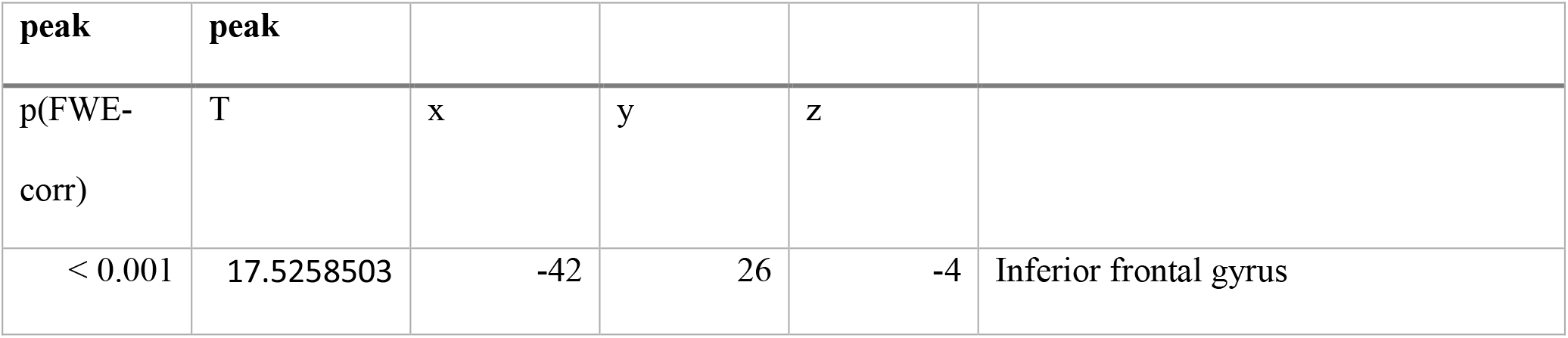
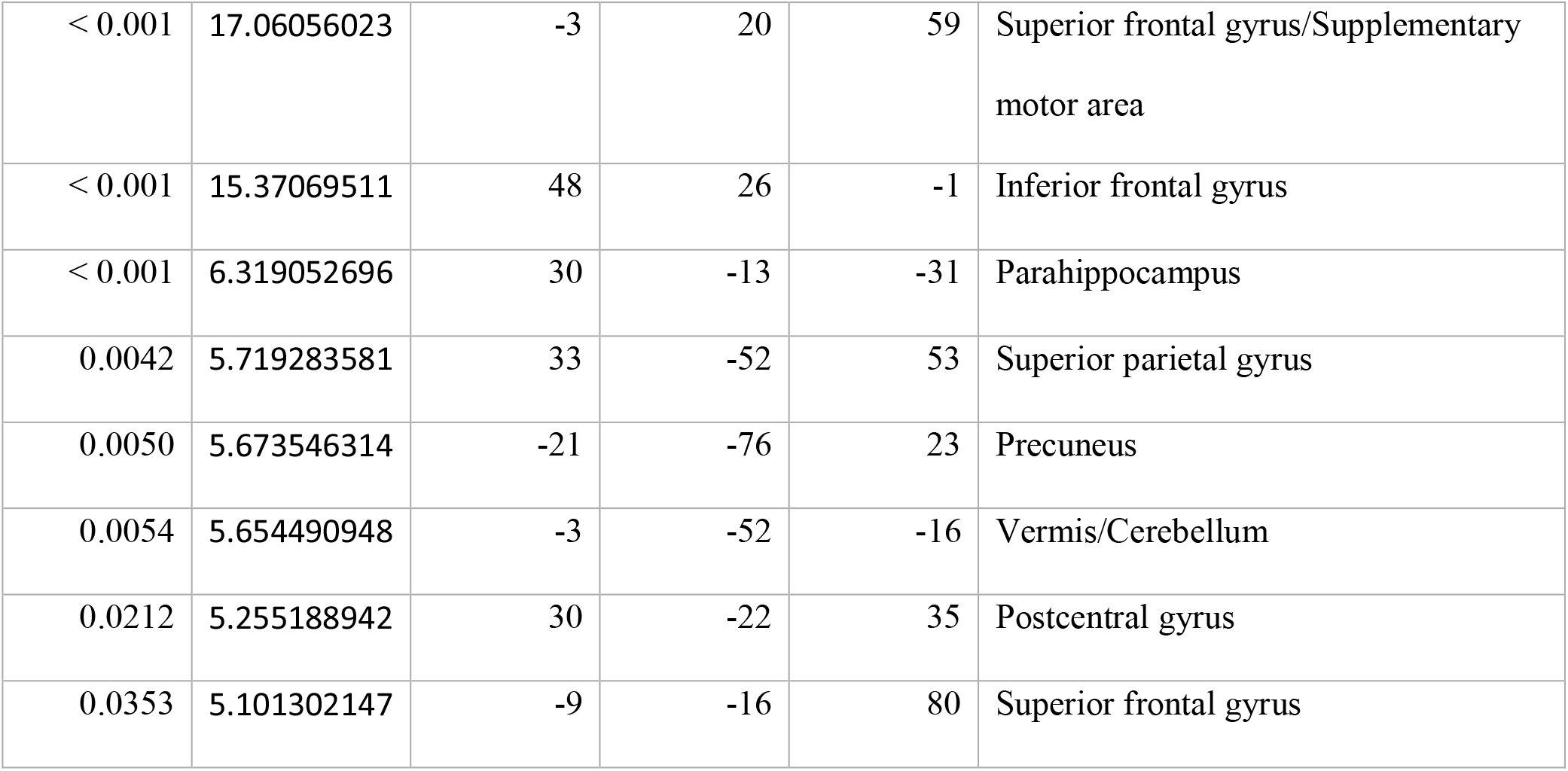
Brain responses to (receiving self compliments > waiting for self compliments)

| peak | peak |  |  |  |  |
| --- | --- | --- | --- | --- | --- |
| p(FWE-corr) | T | x | y | z |  |
| < 0.001 | 17.5258503 | -42 | 26 | -4 | Inferior frontal gyrus |
| < 0.001 | 17.06056023 | -3 | 20 | 59 | Superior frontal gyrus/Supplementary motor area |
| < 0.001 | 15.37069511 | 48 | 26 | -1 | Inferior frontal gyrus |
| < 0.001 | 6.319052696 | 30 | -13 | -31 | Parahippocampus |
| 0.0042 | 5.719283581 | 33 | -52 | 53 | Superior parietal gyrus |
| 0.0050 | 5.673546314 | -21 | -76 | 23 | Precuneus |
| 0.0054 | 5.654490948 | -3 | -52 | -16 | Vermis/Cerebellum |
| 0.0212 | 5.255188942 | 30 | -22 | 35 | Postcentral gyrus |
| 0.0353 | 5.101302147 | -9 | -16 | 80 | Superior frontal gyrus |

**Fig. 1.**
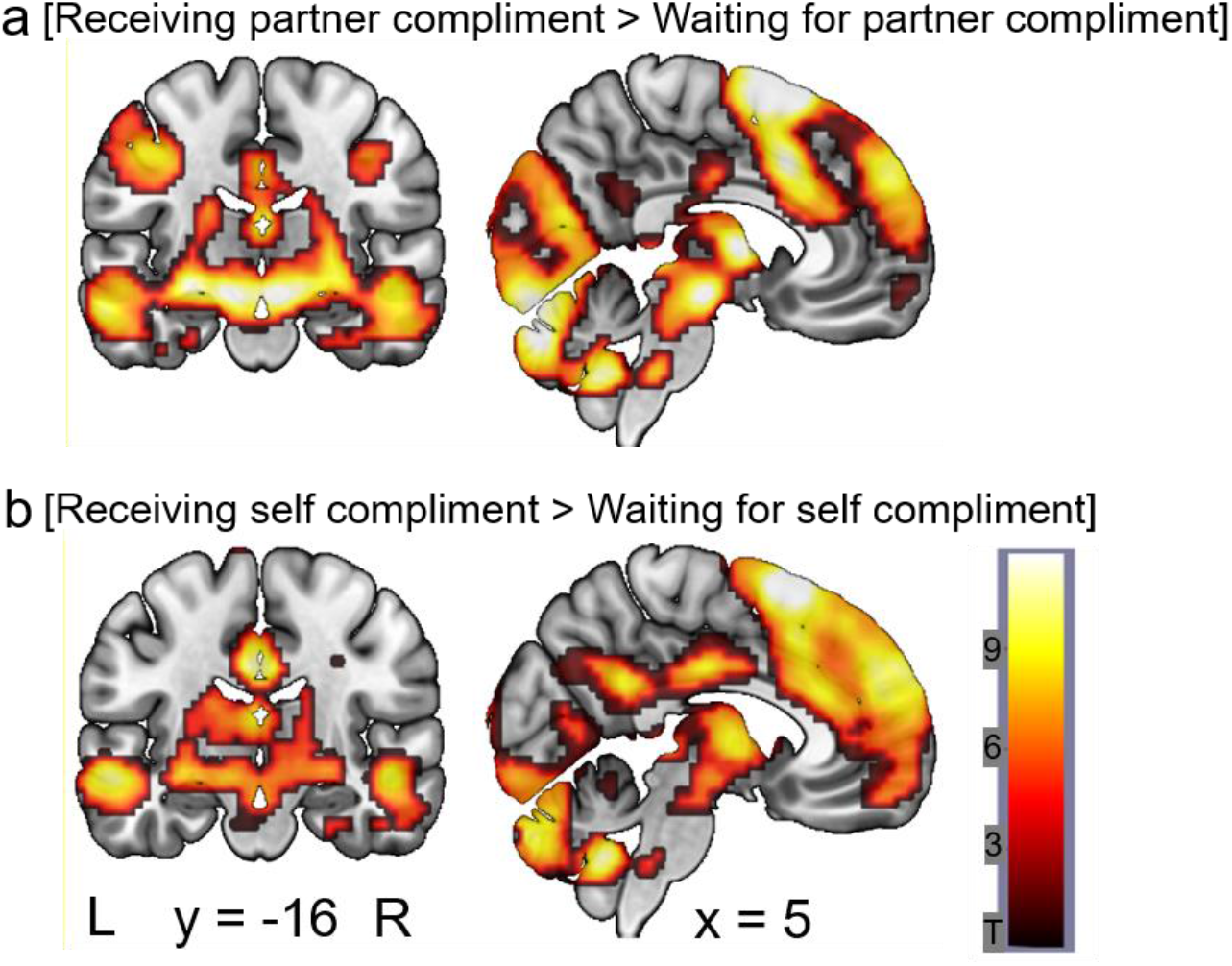
Higher activation during receiving compliments than during waiting, 1a receiving partner compliments 1b receiving self compliments. All figures p<.05, whole brain FWE corrected (x = 5 y= - 16), T-scale applied for both panels

### Activations of individuals when receiving partner vs. self compliments

Contrasting receiving compliments from the partner with self compliments (between the two passive blocks) showed increased VMPFC, ACC, and IFG activity for receiving self compliments (Table 4, Fig 2a) and higher insula, temporal, and amygdala activity when receiving partner compliments. (Table 5, Fig 2b).

**Table 4.** Brain responses to (receiving self compliment > waiting for self compliment) > (receiving partner compliment > waiting for partner compliment)

| peak | peak |  |  |  |  |
| --- | --- | --- | --- | --- | --- |
| p(FWE-corr) | T | x | y | z |  |
| < 0.001 | 7.01081753 | -9 | 26 | -10 | Anterior cingulate |
| < 0.001 | 6.95619154 | -6 | 35 | -22 | Rectal gyrus |
| < 0.001 | 6.37565708 | -21 | 41 | -13 | Middle frontal gyrus |
| < 0.001 | 6.97407198 | 42 | 50 | -4 | Middle frontal gyrus |
| 0.0070 | 5.5989995 | 24 | 53 | -1 | Superior frontal gyrus |
| 0.0002 | 6.51676655 | 54 | -52 | 50 | Inferior parietal lobule |
| 0.0050 | 5.69650269 | 42 | -64 | 53 | Angular gyrus |
| 0.0026 | 5.87921333 | 3 | -28 | 59 | Medial frontal gyrus |
| 0.01062 | 5.47870684 | -9 | -31 | 65 | Medial frontal gyrus |
| 0.0064 | 5.625525 | -36 | -22 | 20 | Insula |
| 0.0070 | 5.59888983 | 36 | -19 | 17 | Insula |
| 0.0098 | 5.50063801 | 42 | 23 | 47 | Middle frontal gyrus |
| 0.0122 | 5.435884 | -9 | 35 | -1 | Anterior cingulate |
| 0.0189 | 5.30584526 | 24 | 41 | -13 | Middle frontal gyrus |
| 0.0251 | 5.21994162 | -42 | -58 | 56 | Partietal inferior<br>gyrus |
| 0.03185 | 5.14777374 | -54 | -55 | 41 | Partietal inferior<br>gyrus |

**Table 5.** Brain responses to (receiving partner compliment > waiting for partner compliment) > (Receiving self compliment > waiting for self compliment)

| **peak** | **peak** |  |  |  |  |
| --- | --- | --- | --- | --- | --- |
| p(FWE-corr) | T | x | y | z |  |
| < 0.001 | 7.21621704 | -30 | -64 | -22 | Cerebellum |
| < 0.001 | 6.71068478 | -42 | -61 | -25 | Cerebellum |
| < 0.001 | 6.43157673 | -12 | -67 | -16 | Cerebellum |
| < 0.001 | 7.18531132 | -48 | -13 | 41 | Pre/postcentral gyrus |
| 0.0072 | 5.59245872 | -48 | -10 | 59 | Precentral gyrus |
| < 0.001 | 6.9873867 | 42 | 17 | -28 | Superior temporal gyrus |
| 0.0012 | 6.0868845 | 48 | 8 | -7 | Insula |
| 0.0140 | 5.39685678 | 36 | -1 | -25 | Amygdala |
| < 0.001 | 6.43445444 | -39 | 11 | -28 | Superior temporal gyrus |
| 0.0011 | 6.10743856 | 45 | -10 | 41 | Precentral gyrus |
| 0.0014 | 6.04848385 | 57 | -4 | 44 | Precentral gyrus |
| 0.0017 | 5.99132061 | 48 | -1 | 59 | Frontal middle gyrus |
| 0.0054 | 5.67191124 | 27 | -7 | -13 | Amygdala |
| 0.0073 | 5.58856153 | -6 | -88 | -10 | Calcarine/Lingual gyrus |
| 0.0093 | 5.51773834 | 3 | 2 | 68 | Superior frontal<br>gyrus/Supplementary motor area |
| 0.0100 | 5.4936924 | -15 | -1 | 80 | Superior frontal gyrus |
| 0.012 | 5.44215488 | 9 | -13 | -13 | Hippocampus |
| 0.0125 | 5.42972374 | -6 | -82 | 17 | Cuneus |
| 0.0197 | 5.29384518 | 3 | -85 | 38 | Precuneus |
| 0.0242 | 5.23110056 | -30 | -13 | -13 | Hippocampus |
| 0.0357 | 5.11234665 | 9 | 14 | 38 | Middle cingulate |
| 0.0458 | 5.03419256 | -30 | -61 | -49 | Cerebellum |
| 0.0469 | 5.02685928 | 15 | -19 | -13 | Hippocampus |

**Fig. 2.**
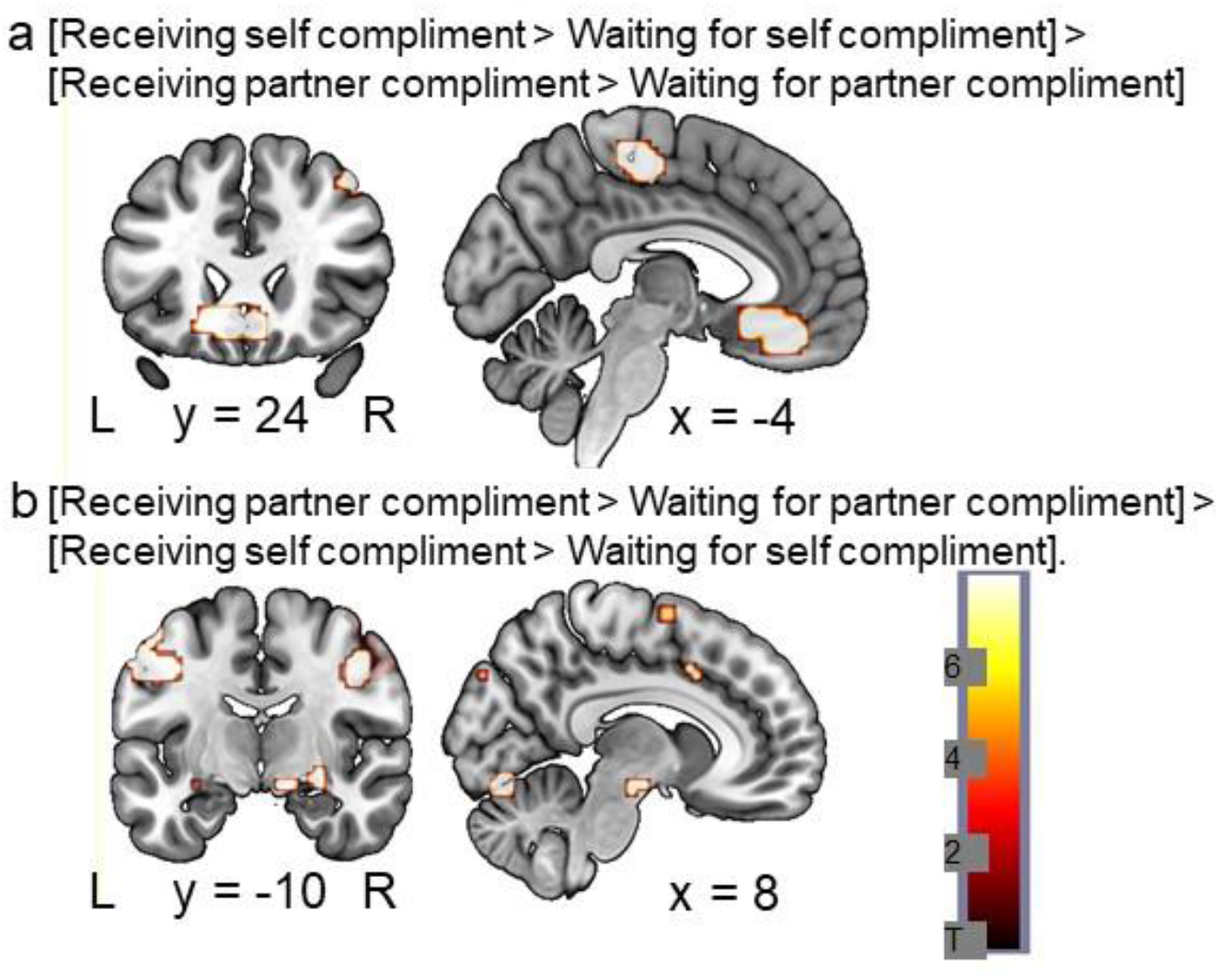
Higher brain activation during partner-than during self compliments, 2a receiving self compliments (receive self-compliments (receive > wait) > receive partner-compliments (receive > wait)) (x =-4, y= 24), 2b receive partner compliments (partner-compliments (receive >wait) > self compliments (receive > wait)) (x= 8, y=-10)

### Activation of individuals when sending compliments

Comparing brain responses during the blocks of sending and receiving of partner compliments, we found that receiving involves larger insula, and hippocampus activity (Table 6, Fig 3a) but selecting/sending compliments for the partner involved an even broader limbic and reward network, including a large cluster around the ventral striatum, TPJ and the cingulate gyrus (Table 7, Fig 3b and for beta estimates for the activity of the ventral striatum see Fig 4).

**Table 6.** Brain responses to (receiving partner’s compliment > waiting for partner’s compliment) > (choosing compliment for partner > observing sending)

| peak | peak |  |  |  |  |
| --- | --- | --- | --- | --- | --- |
| p(FWE-corr) | T | x | y | z |  |
| < 0.001 | 12.1697969 | 21 | -91 | -34 | Cerebellum |
| < 0.001 | 6.47661924 | 3 | -88 | -19 | Cerebellum |
| < 0.001 | 11.8543653 | -33 | -88 | -31 | Superior/Middle occipital gyrus |
| < 0.001 | 11.2321196 | -21 | -79 | -34 | Cerebellum |
| < 0.001 | 10.5057125 | 0 | -1 | 20 | Corpus callosum |
| < 0.001 | 10.3181181 | -12 | -4 | 29 | Middle cingulate |
| < 0.001 | 8.00352383 | -18 | -22 | 29 | Nucleus caudatus/Middle cingulate |
| < 0.001 | 9.35983181 | -54 | -67 | 41 | Angular gyrus |
| < 0.001 | 8.50362015 | -60 | -58 | 29 | Supramarginal gyrus |
| < 0.001 | 8.44472694 | -60 | -46 | 50 | Supramarginal gyrus |
| < 0.001 | 8.84178734 | 57 | -61 | 11 | Superior temporal gyrus |
| < 0.001 | 6.97519064 | 66 | -55 | 20 | Superior temporal gyrus |
| 0.0054 | 5.63413525 | 69 | -40 | 29 | Supramarginal gyrus |
| < 0.001 | 8.6882925 | 6 | 56 | 38 | Superior frontal gyrus |
| < 0.001 | 8.64810467 | 6 | 26 | 68 | Superior frontal gyrus |
| < 0.001 | 7.82007265 | -6 | 50 | 26 | Medial Frontal gyrus |
| < 0.001 | 7.84036493 | -33 | -52 | 2 | Lingual gyrus |
| 0.0014 | 6.01714706 | -24 | -37 | 14 | Hippocampus |
| 0.0025 | 5.84849262 | -18 | -43 | 14 | White matter |
| < 0.001 | 7.54454184 | 27 | -7 | -7 | Globus pallidus/Lentiform nucleus |
| < 0.001 | 7.52939987 | 48 | 2 | -28 | Middle temporal gyrus |
| < 0.001 | 7.01410103 | 48 | 11 | -28 | Temporal pole |
| < 0.001 | 7.29358339 | -45 | 20 | 53 | Middle frontal gyrus |
| < 0.001 | 7.24051571 | -42 | 11 | -34 | Middle frontal gyrus |
| < 0.001 | 7.18364382 | 45 | -10 | 35 | Precentral gyrus |
| < 0.001 | 6.34821653 | 33 | -46 | 2 | Lingual gyrus |
| < 0.001 | 6.16943264 | 3 | -55 | 32 | Middle cingulate/Precuneus |
| 0.0010 | 6.10089874 | -57 | -28 | -19 | Temporal inferior gyrus |
| 0.0023 | 5.8795042 | -63 | -19 | -19 | Temporal middle gyrus |
| 0.0012 | 6.0625968 | 0 | -28 | 17 | Thalamus |
| 0.0020 | 5.90837193 | 57 | -1 | 11 | Insula |
| 0.0028 | 5.81837273 | -42 | -13 | 35 | Postcentral/Precentral gyrus |
| 0.0042 | 5.7014122 | -36 | -13 | 26 | Insula |
| 0.0053 | 5.63772678 | 42 | 5 | -4 | Insula |
| 0.0069 | 5.56073284 | -27 | -1 | -22 | Amygdala |
| 0.0192 | 5.25661325 | 54 | -25 | -13 | Temporal middle |
| 0.0212 | 5.22633171 | 57 | 11 | 2 | Insula |
| 0.0247 | 5.1791029 | 51 | 23 | 5 | Inferior frontal gyrus |
| 0.0339 | 5.08146048 | 57 | 20 | -28 | Middle temporal pole |
| 0.0399 | 5.03027153 | -33 | -7 | -19 | Parahippocampus |
| 0.04829 | 4.96986198 | -54 | -61 | -28 | Cerebellum |

**Table 7.**
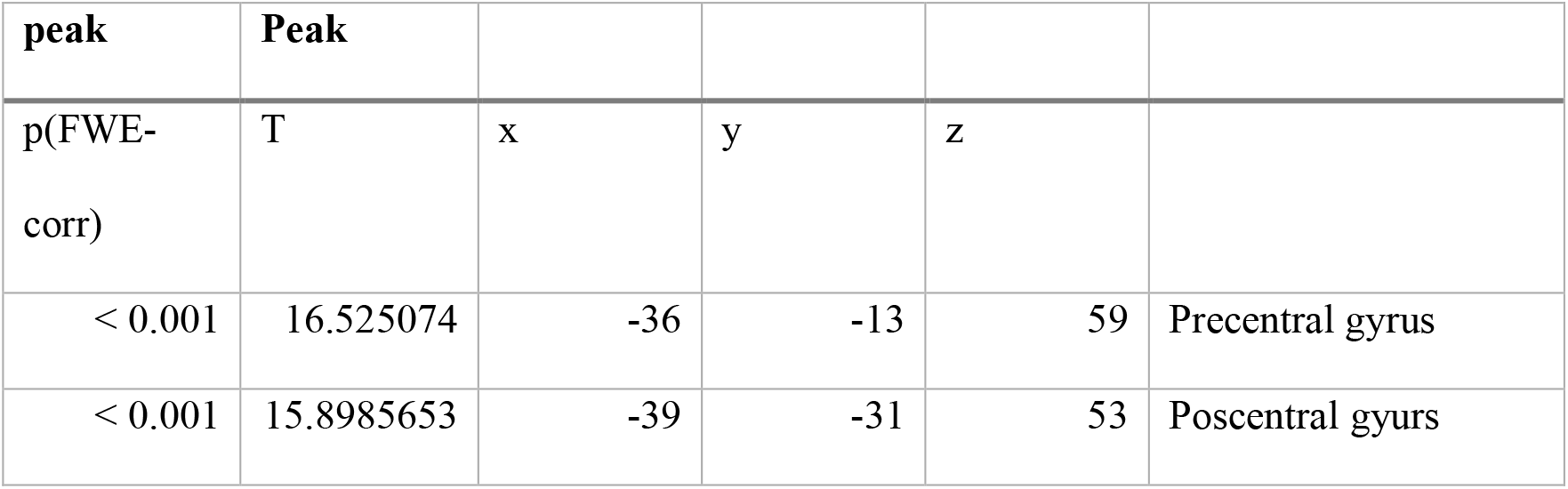
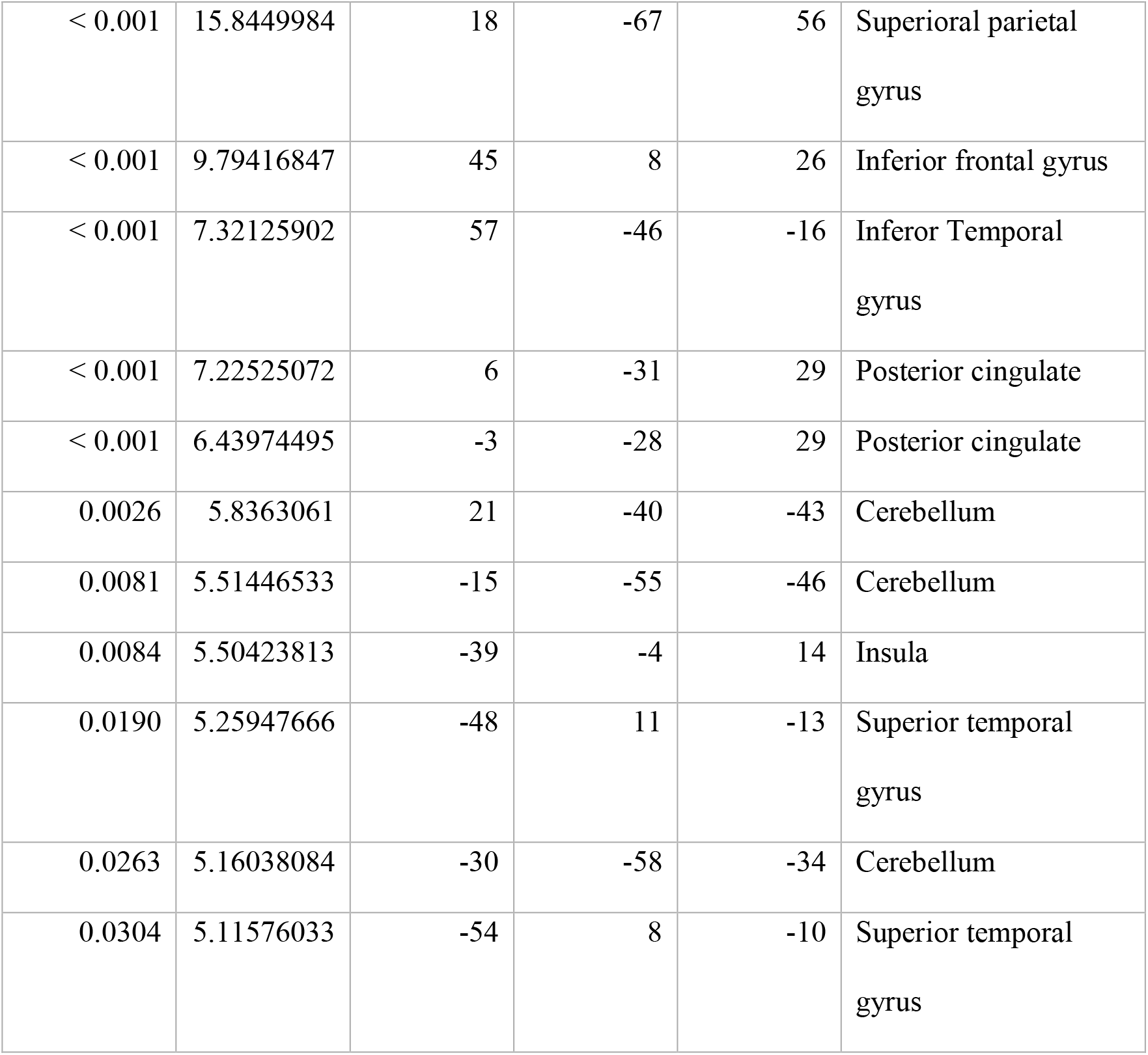
Brain responses to (choosing compliment for partner > observing sending) > (receiving partner’s compliment > waiting for partner’s compliment)

| **peak** | **Peak** |  |  |  |  |
| --- | --- | --- | --- | --- | --- |
| p(FWE-corr) | T | x | y | z |  |
| < 0.001 | 16.525074 | -36 | -13 | 59 | Precentral gyrus |
| < 0.001 | 15.8985653 | -39 | -31 | 53 | Poscentral gyurs |
| < 0.001 | 15.8449984 | 18 | -67 | 56 | Superioral parietal<br>gyrus |
| < 0.001 | 9.79416847 | 45 | 8 | 26 | Inferior frontal gyrus |
| < 0.001 | 7.32125902 | 57 | -46 | -16 | Inferior Temporal<br>gyrus |
| < 0.001 | 7.22525072 | 6 | -31 | 29 | Posterior cingulate |
| < 0.001 | 6.43974495 | -3 | -28 | 29 | Posterior cingulate |
| 0.0026 | 5.8363061 | 21 | -40 | -43 | Cerebellum |
| 0.0081 | 5.51446533 | -15 | -55 | -46 | Cerebellum |
| 0.0084 | 5.50423813 | -39 | -4 | 14 | Insula |
| 0.0190 | 5.25947666 | -48 | 11 | -13 | Superior temporal<br>gyrus |
| 0.0263 | 5.16038084 | -30 | -58 | -34 | Cerebellum |
| 0.0304 | 5.11576033 | -54 | 8 | -10 | Superior temporal<br>gyrus |

**Fig. 3.**
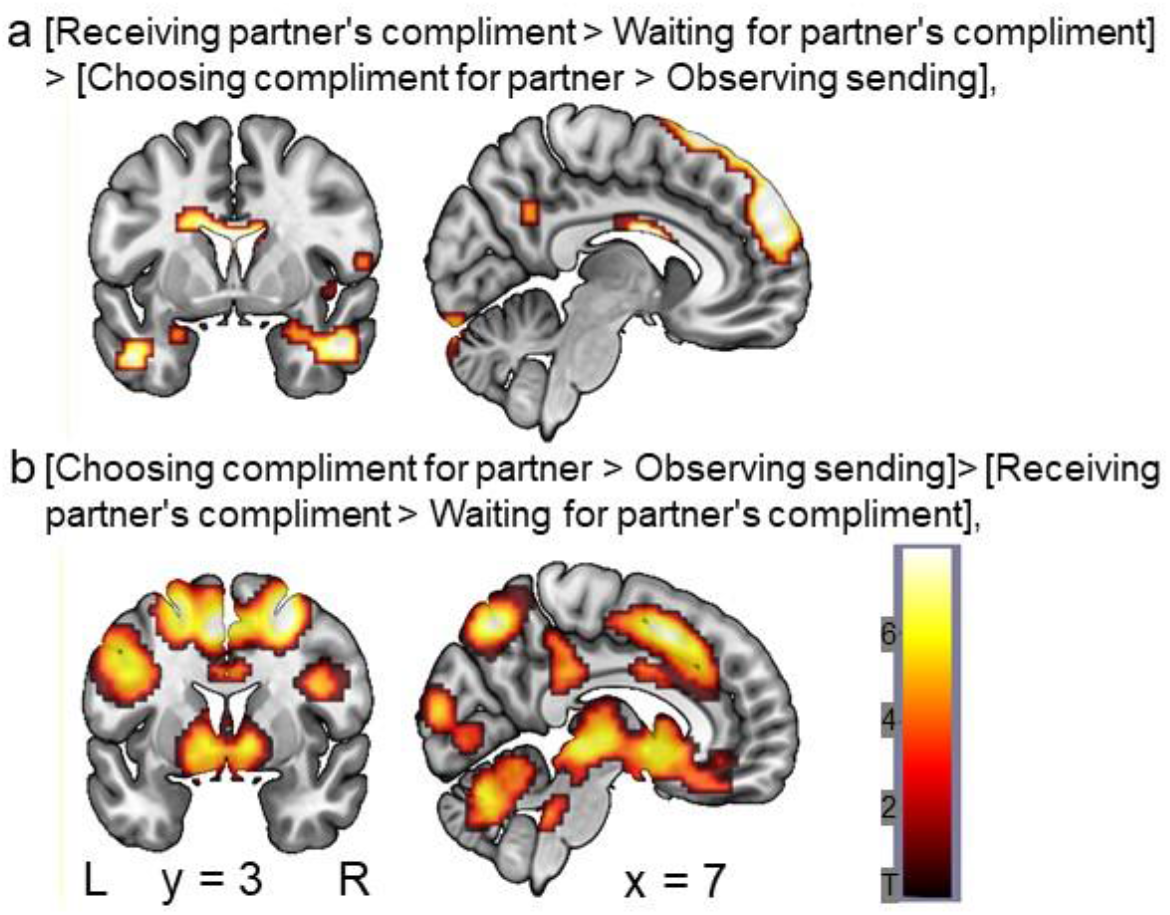
Activation during receiving compared to compliment sending. 3a receiving partner compliment (receive > wait) > sending (choose > observe). 3b sending (choose > observe) > receive partner-compliment (receive > wait), both x 7 = y = 3, t-scale applies to both panels.

**Fig. 4.**
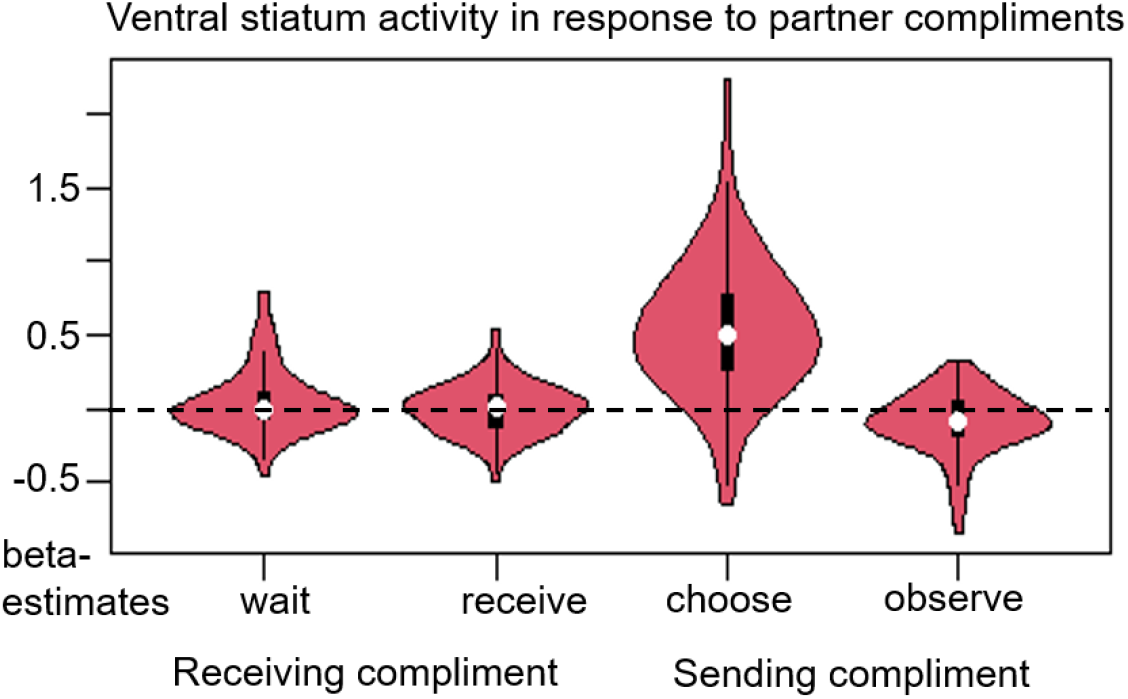
Beta estimates on ventral striatum activation during the experimental phases of sending and receiving partner compliments; white dots indicate means, black bars indicate SEM.

We found no sex differences in any comparison.

Taken together, these results suggest similar activation patterns for self compliment and partner compliment in the paradigm: elevated activation in DL/VMPFC, precuneus, and temporal gyrus when receiving compliments. DLPFC and posterior cingulate are especially sensitive to receiving partner compliments, while temporal lobe and amygdala respond to the anticipation of partner compliments. Interestingly, choosing and sending compliments yielded the strongest activation patterns in the limbic, mentalizing (ToM), and reward systems.

### Results of questionnaire data

A Wilcoxon test (with N = 86, who completed the questionnaires) suggest that participants reported significantly (z = -7.223, p <. 001) more subjective joy during the partner compliment phase (Median = 9, Range 5 – 9) than during the self-compliment phase (Median 6, Range 2 – 9).

## Discussion

For most adult humans, couple relationships are *the* most relevant social relationship, and interacting with the partner modulates momentary affect and long-term health-related outcomes (Braithwaite and Holt-Lunstad, 2017). Exchanging praise and compliments are one element of positive couple interaction and specific compliments in the relationship are assumed to increase social identity (Ellemers, Kortekaas & Ouwerkerk, 1999). The rationale of the present study was to investigate the neural responses when sending and receiving such compliments, as well as receiving self compliments. In summary, we found that both receiving compliments from the partner and self-generated positive attributes activated the salience and limbic networks, as well as the mirror neuron system, as hypothesized. Differential effects occurred especially during the anticipation of the response to a compliment.

The complex activation pattern to receiving compliments corresponded to the activation seen in previous research investigating the reading of emotionally loaded content (Hsu *et al*., 2014). Prefrontal and temporal areas, as well as the insula, were involved in both receiving partner compliments and self compliments. This is in line with the notion that the general processing of self-referential information involves the reward circuitry (Frewen *et al*., 2020) and the dorsal striatum in particular (as part of the nigrostriatal pathway) is involved in comparing predicted and received reward (Oyama *et al*., 2010).

Amygdaloid responses during the anticipation of partner compliments, relate to the ‘emotional’ salience network, but also with social reward (Chan *et al*., 2018). Receiving partner compliments included activation of ACC and temporal gyri. Such activation patterns are part of the “social brain” (Kennedy and Adolphs, 2012) and are involved in successful communication and mentalizing (Van Overwalle and Baetens, 2009; Laurita *et al*., 2017). Here, they might serve as an indicator of ToM and the sender’s mental engagement with choosing a particular compliment.

The compliment choosing phase was associated with complex activation patterns in the senders’ brains which included the dopaminergic reward system. The ventral striatum and neural midline structures showed the strongest activation when choosing a compliment as compared to the other conditions (see Fig 4). While this was not hypothesized, these results are well in line with previous reports indicating that emotion sharing might be rewarding (Wagner *et al*., 2014), and can be associated with striatal activation during the anticipation of reward (Filimon *et al*., 2020). Since there was no other experimental condition including non-emotional decision making to compare this data with, we have to interpret this finding with caution, though. Other examples of rewarding anticipation of prosociality include supporting financially family members, which elicits activation in the mesolimbic dopaminergic system (Telzer *et al*., 2010), as well as deciding to donate to charities, which recruited the ventral and dorsal striatum and VTA (Moll *et al*., 2006). Similarly, Harbaugh *et al*. (2007) found that both mandatory and voluntary contributions to charities recruited the same areas. Finally, Izuma *et al*. (2010) reported that ventral striatum activity to charitable donations increased in the presence of others, suggesting that this region may be particularly sensitive to social rewards. Our present results add to this line of literature by showing for the first time the differential contributions of dorsal striatum to receiving a treat oneself and of ventral striatum to selecting a treat for someone else during live social interaction.

Our data imply that throughout all conditions, the senders paid close attention to the reaction of their partners during compliment sharing: Activation in oculo-, pre- and motor areas, as well as areas associated with showing emotional, mostly happy, faces such as pSTS and DMPFC suggest involvement of the emotionally ‘extended mirror neuron network’ (Van der Gaag *et al*., 2007). Positive affect and frontal activity during emotional partner interaction were also reported in a recent study using EEG (Packheiser *et al*., 2021).

In summary, by using a somewhat naturalistic interaction paradigm, the present study design builds on previous research on reward-related brain activation in romantic couples such seeing as pictures from the partner (Acevedo *et al*., 2012) and extends existing data to a more dynamic couple interaction. To our knowledge, this work is the first to investigate the neural underpinnings of positive emotional interaction between romantic couples using individually meaningful attributes characterizing the relationship and the participants involved, namely self-generated compliments.

The specific areas found to be involved in couple’s compliment sharing are known for social cognition processes, social reward processing, ToM, and facial mimicry (Jabbi and Keysers, 2008; Kennedy and Adolphs, 2012). The involvement of the dopaminergic reward system in particular might serve as an important neurobiological mechanism underlying the ever rewarding aspects of lasting couple relationships. Interestingly, these brain areas are also involved in the action of neuropeptides promoting social behavior, such as oxytocin (Riem *et al*., 2012; Kreuder *et al*., 2018). Oxytocin has been shown to interact with the reward system, for example when study participants observed the face of their romantic partner (Scheele *et al*., 2013), and also to influence the appraisal of the relationship (Aguilar-Raab *et al*., 2019). Furthermore, oxytocin is known to promote health-beneficial effects such as regulation of the stress axes during couple interaction (Ditzen *et al*., 2009; Zietlow *et al*., 2018). Therefore, the neural networks reported here and the role of oxytocin might provide a potential neurobiological pathway underlying the association of couple relationships and health.

Our study has strengths but also some limitations. Investigating heterosexual romantic couples only and having them name, choose, and send the compliments helped create an individualized interaction scenario. The paradigm comprised receiving unknown compliments from the partner and known self-compliments while always seeing the partner via video transmission as part of a naturalistic social exchange. Therefore, we can neither rule out that the found differences between partner- and self-compliment are due to novelty nor that some kind of interaction has taken place during the self-compliment phase via facial expressions. Other aspects that differed between task phases were the active or passive role of leading the interaction or making decisions in general.

The selected heterosexual monogamous sample allows no extrapolation to unacquainted individuals, platonic friend dyads or same-sex couples, though. Furthermore, the sample consisted of healthy young couples reporting high relationship satisfaction only. Given inconsistent effects of instructed partnership appreciation in clinical samples (Warth *et al*., 2020) or couples in therapy (Aguilar-Raab *et al*., 2018), we cannot extrapolate our findings to marital problems or patient populations (see for instance a study in couple with substance abuse by Flanagan *et al*., 2018). On the other hand, our findings may still be applicable for some cultures or couple circumstances, since our participants came from Europe and North Africa (15 different nations and 12 mother tongues) therefore generalizability to those parts of the world is given and the individualized compliments have accounted for potential differences. Future studies could systematically investigate cultures and contexts, clinical samples, and couples in the LGBTQIA+ spectrum. We assume similar basic neural effects in all couples though.

In conclusion, our data show substantial involvement of limbic structures during instructed yet individualized couples compliment sharing. The involvement of dopaminergic areas, is evident not only when receiving compliments but is strongest in the ventral striatum when selecting compliments for the partner. This suggests a role of neural reward processes when giving a treat to the loved one - which might contribute to the maintenance of lasting relationships beyond the mere receipt of affection and support.

## AUTHORS’ CONTRIBUTION STATEMENT

B.D., P.K., G.S. E.B and M.E. designed the study; G.S. and M.E. lead the study; G.S., E.B. and M.E. collected the data; M.F.G. and E.B. established the experimental set-up; G.S. M.F.G and M.E. ran the reported analyses, M.E., and B.D. wrote the manuscript; All authors provided comments on the manuscript.

## ACKNOWLEDGMENTS

We wish to thank Fabienne Ibert, Jasmin Buchholz, Julia Hein and Laura Kampouridis for their skilled support in data collection and Sarah Fancy for proofreading the manuscript.

